# Engineering a perfusion bioreactor system for hiPSC-derived progenitor co-culture capturing microglial features in CNS development

**DOI:** 10.1101/2025.07.08.663740

**Authors:** Catarina M. Gomes, Inês de Sá, Margarida Delgado, Paula M. Alves, Catarina Brito

## Abstract

Microglia are critical regulators of brain homeostasis and immune responses in the central nervous system (CNS). However, existing human-based models fail to reproduce the early and complex microglia-neural cell interactions. The differentiation of human induced pluripotent stem cells (hiPSCs) into specialized cell types offers promising avenues for understanding human development and disease modeling. Herein, we explore the differentiation of hiPSC-derived erythromyeloid progenitors (iEMPs) and their 3D co-culture with hiPSC-derived neurospheres, utilizing the Ambr® 250 Modular system. The aim of this research was to build a complex co-culture model between iEMP and neurospheres in a scalable and controlled environment. Our results demonstrate that the Ambr® 250 Modular system effectively supports the co-culture process, with iEMPs integration into the neurospheres, exhibiting cell density, aggregate morphology and concentration similar to the neurosphere monocultures. The co-culture environment induced the upregulation of transcription factors critical for microglial lineage commitment. iEMP-neurospheres displayed a unique secretory profile, releasing proteins involved in extracellular matrix remodeling and neuronal differentiation, essential for microenvironment remodeling. In conclusion, this study underscores the role of iEMPs in CNS development and presents a robust platform for preclinical research.

## Introduction

Microglia, the resident immune cells of the central nervous system (CNS), are pivotal in orchestrating neurodevelopmental processes and maintaining tissue homeostasis. They are actively involved in phagocytosing apoptotic cells, pruning synapses, and modulating neurogenesis, all essential for brain development (Su et al., 2014; Yamamiya et al., 2019). Microglia dysfunction during development has been increasingly implicated in various neurodevelopmental disorders, including autism spectrum disorders (ASD) (Harrington et al., 2020; Xu et al., 2020b), schizophrenia (Sellgren et al., 2019) and Rett syndrome (Maezawa and Jin, 2010; Cao et al., 2024). Animal models have been instrumental in preclinical research, enriching our knowledge of cellular and molecular mechanisms of brain development, ageing, and neurological diseases (Gao et al., 2023; Yin et al., 2023). However, significant differences between human and animal models limit our interpretation and translation of generated data. The availability of functional human brain tissue from healthy and diseased individuals is limited, especially tissue from rare disorders. The emergence of human induced pluripotent stem cells (hiPSCs) provides new opportunities to overcome these challenges (Avior et al., 2016; Cerneckis et al., 2024)}. By focusing on neural differentiation of hiPSCs, in vitro two-dimensional (2D) human neural cell culture and three-dimensional (3D) neurospheroid (iNSpheroid) and organoid models have helped uncover mechanisms underlying human brain development and diseases (Qian et al., 2016a, 2019; Kelley and Pa□ca, 2022). Given the central role of microglia in neurodevelopmental disorders, incorporating them into three-dimensional (3D) in vitro models is essential for studying disease mechanisms and testing potential therapeutics. Advancing these models will be crucial for developing targeted therapies for neurodevelopmental disorders, enabling a deeper understanding of microglial contributions to brain development and disease. However, hiPSC-derived CNS cells, such as astrocytes and microglia, often show limited cellular and functional maturation and heterogeneity either cultured in 2D or 3D (Cerneckis et al., 2024).

hiPSC-derived microglia have been explored for studying in vitro their development and function in human-specific pathological contexts (Muffat et al., 2016; Abud et al., 2017; Amos et al., 2017; Douvaras et al., 2017; Haenseler et al., 2017; Pandya et al., 2017; McQuade et al., 2018; Xu et al., 2020a; Banerjee et al., 2020; Trudler et al., 2021; Funes and Bosco, 2022; Lanfer et al., 2022). However, traditional 2D culture systems fail to capture the complex cellular interactions and the human brain microenvironments, leading to significant differences in microglial morphology and gene and protein expression compared to in vivo conditions (Gosselin et al., 2017; Vahsen et al., 2022). In contrast, iNSpheroid and brain organoids provide higher physiological relevance, recapitulating the cellular heterogeneity and interactions occurring in the brain (Bayó-Puxan et al., 2018; Simão et al., 2018; Gomes et al., 2024)(reviewed in (Del Dosso et al., 2020)). Despite these advantages, these models still exhibit key limitations, including the lack of vascularization, immature cellular phenotypes, variability in differentiation protocols, and often lack a functional immune system, particularly microglia, which play a crucial role in neurodevelopment, synaptic remodeling, and neuroinflammatory responses (Zhao et al., 2021; Ottaviani et al., 2023; Werschler et al., 2024; Di Stefano et al., 2025). Although the advent of microglia-containing 3D models represents a significant advancement, several challenges remain, including microglial heterogeneity and inflammatory profile, their long-term survival and functional maintenance, and scalability issues that limit the throughput necessary for preclinical applications (reviewed in (Dumas et al., 2021; Di Stefano et al., 2025)).

To further advance preclinical research, there is an emerging need to develop immunocompetent platforms that integrate human microglia with other neural cell types in a scalable and reproducible manner. Stirred-tank bioreactors have demonstrated significant potential in addressing these needs by offering precise control over environmental parameters such as pH and oxygen control, which are critical for maintaining the health and functionality of hiPSC-derived cells (Simão et al., 2016, 2018; Bayó-Puxan et al., 2018; Ho et al., 2022). Indeed, in recent years oxygen concentration has been shown to be one of the key factors influencing human pluripotent stem cell (hPSC) characteristics, differentiation and function (Iida et al., 2013; Kusuma et al., 2014; Sugimoto et al., 2018; Zhi et al., 2023; Vicente et al., 2024, 2025). We have previously reported the 3D human iNSpheroid model, generated by 3D differentiation of hiPSC-derived neural progenitor cells (iNPCs) in stirred tank bioreactors. This two-stage methodology allows the generation and maintenance of neurospheres, consisting of a uniform population of iNSCs, and subsequent co-differentiation of neurons, astrocytes, and oligodendrocytes from these structures (iNSpheroids). We have demonstrated that the cells in the iNSpheroids can produce and deposit brain-like extracellular matrix (ECM), fostering cell-cell and cell-ECM interactions that recapitulate brain cell architecture features and pathological processes (Bayó-Puxan et al., 2018; Simão et al., 2018).

Herein, we introduced a novel methodology to co-culture hiPSC-matched microglia precursor cells (erythromyeloid progenitors, iEMPs) and iNPCS towards the development of an isogenic immunocompetent platform to investigate the poorly understood relationship between the progenies of these cells. We leveraged the Ambr® 250 Modular, a high-throughput and scalable system, making it ideally suited for the large-scale production of complex 3D models such as iNSpheroids and organoids. An in-house developed perfusion system (Simão et al., 2016), was adapted to the Ambr® 250 Modular, given the absence of commercially available solutions that can accommodate 3D culture needs. This approach models microglia-neural progeny interactions in a scalable system. It offers valuable insights into microglial roles in neurodevelopment and neurological disorders, improving preclinical models for drug discovery and therapy testing.

## Materials and Methods

### Human induced pluripotent stem cell (hiPSC) culture

Four hiPSC lines were used in this work, as described in table 1. The hiPSC were expanded on Matrigel® hESC-Qualified Matrix, LDEV-free (Corning®), in mTeSR™1 cGMP medium (StemCell Technologies) under feeder-free culture conditions. A complete medium exchange was performed every day. Cells were maintained under a humidified atmosphere with 5 % CO_2_, at 37 °C.

**Table 1:**
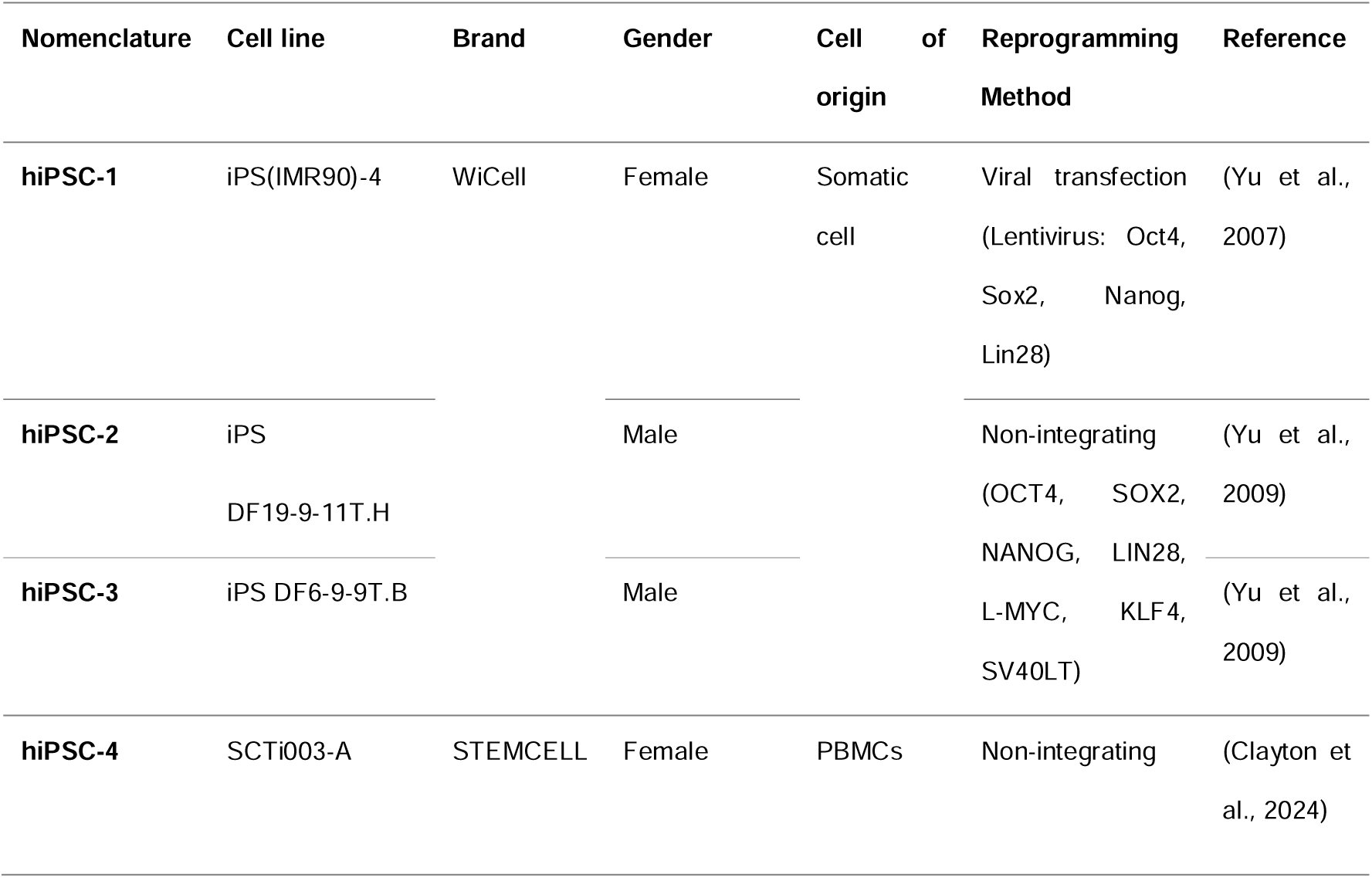
hiPSC lines nomenclature and origin.

### Generation of iNPCs

iPSC(IMR90)-clone 4 and iPSC DF6-9-9T.B cells were plated at 0.8×10^5^ cell/cm^2^ in mTeSR™1 cGMP medium and allowed to adhere to pre-coated Matrigel® hESC-Qualified Matrix-coated plates, for 24 hours. Afterward, cells were cultured in either (i) DualSMADi culture medium (iPSC(IMR90)-clone 4): N2B27 medium supplemented with SB431542 at 10 μM and LDN193184 at 100 nM (both from STEMCELL™Technologies); or (ii) STEMdiff™ Neural Progenitor Medium (STEMCELL™Technologies) (DF6-9-9T.B). N2B27 medium is composed by ½ DMEM/F-12 and ½ Neurobasal, N2-supplement (1x), B27 supplement without vitamin A (0.5x), GlutaMAX (1x), Penicillin-Streptomycin (1%), MEM Non-Essential Amino Acids Solution (1%) and 2-mercaptoethanol (50 μM) (all from Gibco™), and Insulin (20 μg/mL) (Sigma-Aldrich). A 100% medium exchange was performed daily, for 7 (DF6-9-9T.B) or 10 days (iPSC(IMR90)- clone 4). By the end of differentiation, cells were recovered and plated on poly-L-ornithine-laminin (PLOL)-coated surfaces. PLOL coating was prepared by performing a 3-hour incubation at 37°C with 0.16 mg/mL solution of poly-L-ornithine in DPBS (with Ca^2+^ and Mg^2+^), followed by a washing step and a 3-hour incubation at 37°C with 1 μg/mL solution of laminin in DPBS (with Ca^2+^ and Mg^2+^). Cells were maintained and expanded on PLOL-coated surfaces in a NPC expansion medium (EM), composed of DMEM/F12 media with Glutamax (Life Technologies) supplemented with 1% N2 supplement (Life Technologies), 0.1% B27 supplement (Life Technologies), 1.6 μg/mL glucose (Sigma-Aldrich), 20 μg/mL insulin (Sigma-Aldrich), 20 ng/mL rhubFGF (Peprotech) and 20 ng/ml rhu-EGF (Sigma-Aldrich) (Koch et al., 2009). The medium was changed every other day until confluent cell culture was obtained.

NPCs were passaged at 90% confluence (typically every 3-4 days). Cells were dislodged by 0.05% Trypsin-EDTA (1-2 min at RT), which was neutralized with DMEM supplemented with 10% FBS (Life Technologies). Cells were sedimented by centrifugation, resuspended in EM and plated on PLOL-coated T-flasks, at a cell density of 3×10^4^ cell/cm^2^. A 50% media exchange was performed on day 2 of culture. Cells were maintained under a humidified atmosphere, in a multi-gas cell incubator (Sanyo), with 5% CO_2_ and 3% O_2_, at 37 °C.

### hiPSC-EMP (iEMP) differentiation

hiPSC-1-4 lines were differentiated into EMPs by using the STEMdiff™ Hematopoietic Kit (STEMCELL™ Technologies). One day before induction of differentiation, hiPSCs were harvested through incubation with ReLeSR™ (STEMCELL™ Technologies), according to the manufacturer’s instructions. Cells were seeded in 6-well plates as small colonies (100 – 200 μm in diameter), at 4-10 colonies/cm^2^, in mTeSR™1 supplemented with 10 μM of Y-27632. On the following day, the cell culture medium was replaced with Medium A (STEMdiff™ Hematopoietic Basal Medium containing 0.5% (v/v) of Supplement A) to induce mesoderm differentiation (day 0). On day 2, half of the medium was changed with fresh Medium A. On the following day, the medium was completely exchanged to Medium B (STEMdiff™ Hematopoietic Basal Medium containing 0.5% (v/v) of Supplement B). Half medium exchanges were performed on days 5 and 7 and 10 to promote further lineage commitment towards erythromyeloid cells. By day 12, iEMPs could be harvested from the culture supernatant and frozen at 1-2 million cells per mL in BamBanker ® (LYMPHOTEC Inc.).

### iNSC and iEMP culture in the Ambr® 250 Modular system

iNSCs were expanded in 2D and harvested as described previously (Simão et al., 2016, 2018). The iNSC cell suspension was passed through a 70 µm nylon strainer (Millipore) and diluted to a cell density of 4×10^5^ cell/mL in NPC aggregation medium (AM). AM is formulated similarly to EM except for the reduced concentration of EGF and bFGF (5 ng/mL) and supplementation with ROCK inhibitor (10 μM, Y-27632). Cells were inoculated into a benchtop software-controlled stirred-tank bioreactor (STB) system for parallel cell culture, the Ambr® 250 Modular system (Sartorius Stedim). The single-use bioreactor vessels selected were unbaffled and had a single large elephant ear impeller, providing the gentle mixing needed for 3D cell culture application (Rotondi et al., 2021). Culture conditions were set to maintain cells under 3% dissolved oxygen (15 % of air with 21 % of oxygen, headspace), pH 7.4, 37 °C, and a stirring rate varied according to time of culture, between 160-230 rpm, in down-stirring mode. A 0.5% N_2_ gas flow was added by the sparger to hinder the backflow through this tubbing. After 2 days of iNSC culture, a single iEMP cell suspension was diluted in co-culture medium (CAM) for a starting ratio of 10 iNPC to 1 iEMP cells. Co-culture medium was composed of AM supplemented with interleukin (IL)-3 and M-CSF (10 ng/mL, Preprotech).

### Perfusion system adaptation to Ambr® 250 Modular

After 3 days of culture, perfusion operation mode was activated, with a dilution rate of 0.33 day^-1^ under syringe (inlet) and bi-directional (outlet) pump control. The reservoirs coupled to the STB vessel were used to add fresh medium, with temperature control that ranges from 4-8 °C. As a cell retention device, a stainless-steel frit sparger was connected to the bi-directional pump on each Ambr® 250 Modular vessel slot, using silicon and metal tubing with ¼” OD and 1/8” ID. The stainless-steel frit sparger pore size ranged from 10 to 20 µm, allowing for the simultaneous retention of cell aggregates inside the STB and removal of culture medium, single cells, and cell debris. Perfusion was controlled using automated gravimetric control as described before (Simão et al., 2016).

### Cell concentration and viability

Cell concentration and viability were analysed using the NucleoCounter® NC-200™, a cell counter that uses low-magnification fluorescence microscopy and automated image analysis to identify live and dead cells. For 2D culture systems, these parameters were evaluated through the ‘Viability and Cell Count’ protocol, available in the NucleoView NC-200™ software. For 3D culture, reagents A100 and B were used to lyse the cells and perform a nuclei concentration assessment, using the ‘Viability and Cell Count – A100 and B Assay’ protocol. Samples were prepared and loaded into the Via1-Cassette™ system according to the manufacturer’s instructions for each protocol, and the parameters of interest were further determined.

Cell population doubling time (PDT) was calculated using the following formula:

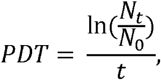

where *N_0_* is the initial number of cells seeded and *N_t_* is the number of cells counted at time *t*.

### Aggregate viability, concentration, and size determination

Aggregate viability was assessed using a combination of fluorescein diacetate (FDA) and propidium iodide (PI). For this, the aggregates were incubated with 1:500 of FDA (20 μg/mL, Sigma-Aldrich™) and 1:1000 of PI (10 μg/mL, Sigma-Aldrich™) in Dulbecco’s phosphate-buffered saline without Mg2+ and Ca2+ (DPBS (-/-), Gibco™) and visualized using a fluorescence microscope (DMI6000, Leica). Images were acquired with a monochrome digital camera (Leica DFC360 FX) and processed using ImageJ software (Schindelin et al., 2012). Aggregate concentration was assessed by manual counting using a phase-contrast microscope. Aggregate size was determined through the analysis of FDA fluorescence images with ImageJ. The area occupied by the aggregates was defined through manual threshold adjustment, and the Feret diameter (in µm) was measured.

### Quantification of extracellular metabolite concentration

Concentrations of glucose (GLC), alanine-glutamine (AQB), glutamine (GLN), pyruvate (PYR), lactate (LAC), ammonia (NH3) and lactate dehydrogenase (LDH) in cell culture supernatants were measured using Cedex Bio Analyzer (Roche). The absolute metabolite concentration (qMet, expressed in pmol/cell) was calculated using the following equation:

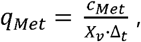

where cMet represents the variation in metabolite concentration with a specific cell concentration average, Xv.

### Click-IT™ EdU assay

Neurospheres with or without iEMP were harvested from the STB vessel and plated on PLOL or Poly-D-Lysine (Gibco™)-coated glass coverslips and incubated, at 37 °C with 5% (v/v) CO_2_ for 24 hours, with 5-ethynyl-2’-deoxyuridine (EdU), at a ratio of 1:1000. After this period, neurospheres were washed with 1% (v/v) bovine serum albumin (BSA, Sigma-Aldrich) in DPBS. Samples were then fixed, for 20 minutes, at room temperature (RT), in 4% (v/v) paraformaldehyde (PFA, Fluka®) and 4% (v/v) sucrose (Sigma-Aldrich®) in DPBS with Mg2+ and Ca2+ (DPBS (+/+), Gibco™). Afterwards, neurospheres were washed twice with 1% (v/v) BSA and incubated for 30 minutes at RT with component E from Click-iT™ EdU Alexa Fluor™ 488 Flow Cytometry Assay Kit (Invitrogen™), diluted in 1% (v/v) BSA. Then, the supernatant was discarded and Click-iT™ reaction cocktail was added, according to the manufacturer’s instructions, and incubated for 1 hour, at RT, in the dark. Two washing steps were performed in component E diluted in 1% (v/v) BSA, iEMPs were further stained with anti-CD45 conjugated antibodies (BioLegend) for 45 minutes at RT. Two additional washing steps were performed and cell nuclei were counterstained using DAPI (4’,6-diamidino-2-phenylindole dihydrochloride, 10 μg/mL, Invitrogen™) diluted 1:1000 in DPBS (+/+). Neurospheres were mounted in ProLong™ Gold Antifade reagent (Invitrogen™) between a microscope slide and a glass coverslip. Preparations were visualized and acquired on a Zeiss LSM 880 point scanning confocal microscope controlled with the Zeiss Zen 2.3 (black edition) software. The obtained images were further processed using the ImageJ software.

### Immunofluorescence microscopy

Neurospheres with or without iEMP were harvested from the STB vessel, washed three times in DPBS (+/+) and fixed in 4% PFA + 4% sucrose in DPBS for 20 min at RT. Afterwards, the neurosphere suspension was washed three times with DPBS, to remove the fixation solution. For immunostaining, block and permeabilization were performed simultaneously with 0.2% fish skin gelatine (FSG) and 0.5% Triton X-100 in DPBS, for 20 min, at RT. Primary antibodies were then incubated for 2 hours at RT and overnight at 4 °C, diluted in 0.2% FSG + 0.5% TritonX-100 in PBS. Neurospheres were washed three times with DPBS and incubated for 2 hours with secondary antibodies diluted in the same solution. Primary and secondary antibodies were used as follows: anti-nestin (1:200, AB5922, Merk Millipore), anti-MAP2 (1:200, AB92434, Abcam), anti-AML1 (1:100, 1674336S, Cell Signaling Technology), anti-PU.1 (1:100, 16789136S, Cell Signaling Technology), IRF-8 (1:100, 16798344S, Cell Signaling Technology), anti-TMEM119 (1:100, 16741134S, Cell Signaling Technology), Alexa Fluor 488 goat anti-mouse IgG, Alexa Fluor 555 donkey anti-rabbit IgG, and Alexa Fluor 647 goat anti-chicken IgY (1:500, A-11001, A-31572, A-32933, Invitrogen). Cell nuclei were counterstained with DAPI diluted in DPBS for 5 min (1:1000, Life Technologies), after which coverslips were washed three times in DPBS. Coverslips were mounted in ProLong™ Gold Antifade Mountant (Life Technologies). Images were acquired using a Leica MICA (Microhub) microscope in confocal mode. The system was controlled using Leica’s LAS X software. Images were processed using ImageJ software and only linear manipulations were performed (Schindelin et al., 2012).

### RT-qPCR

Total RNA was extracted with High Pure RNA Isolation Kit (Roche) or RNeasy Mini Kit (Qiagen, Valencia, CA, USA), according to the manufacturer’s instructions. RNA was quantified in a NanoDrop 2000c (Thermo Scientific) and used for cDNA synthesis. Reverse transcription was performed using High Fidelity cDNA Synthesis Kit (Roche), using Anchored-oligo(dT)18 Primer (Roche), or with the Sensiscript RT Kit (Qiagen), for low cDNA abundant samples, using a mix between Anchored-oligo(dT)18 Primer and random hexamers (Roche). qPCRs were performed in triplicates using the LightCycler 480 SYBR Green I Master Kit (Roche) and the primers listed on Table 2. The reactions were performed with LightCycler 480 Instrument II 384-well block (Roche). Quantification cycle values (Cq’s) and melting curves were determined using the LightCycler 480 Software version 1.5 (Roche). All data were analysed using the 2-ΔΔCt method for relative gene expression analysis (Livak and Schmittgen, 2001). Changes in gene expression were normalized using the housekeeping genes RPL22 (ribosomal protein L22), GADPH (Glyceraldehyde 3 Phosphate Dehydrogenase) as internal controls. Statistical analysis was carried out using GraphPad Prism 9 software.

**Table 2:**
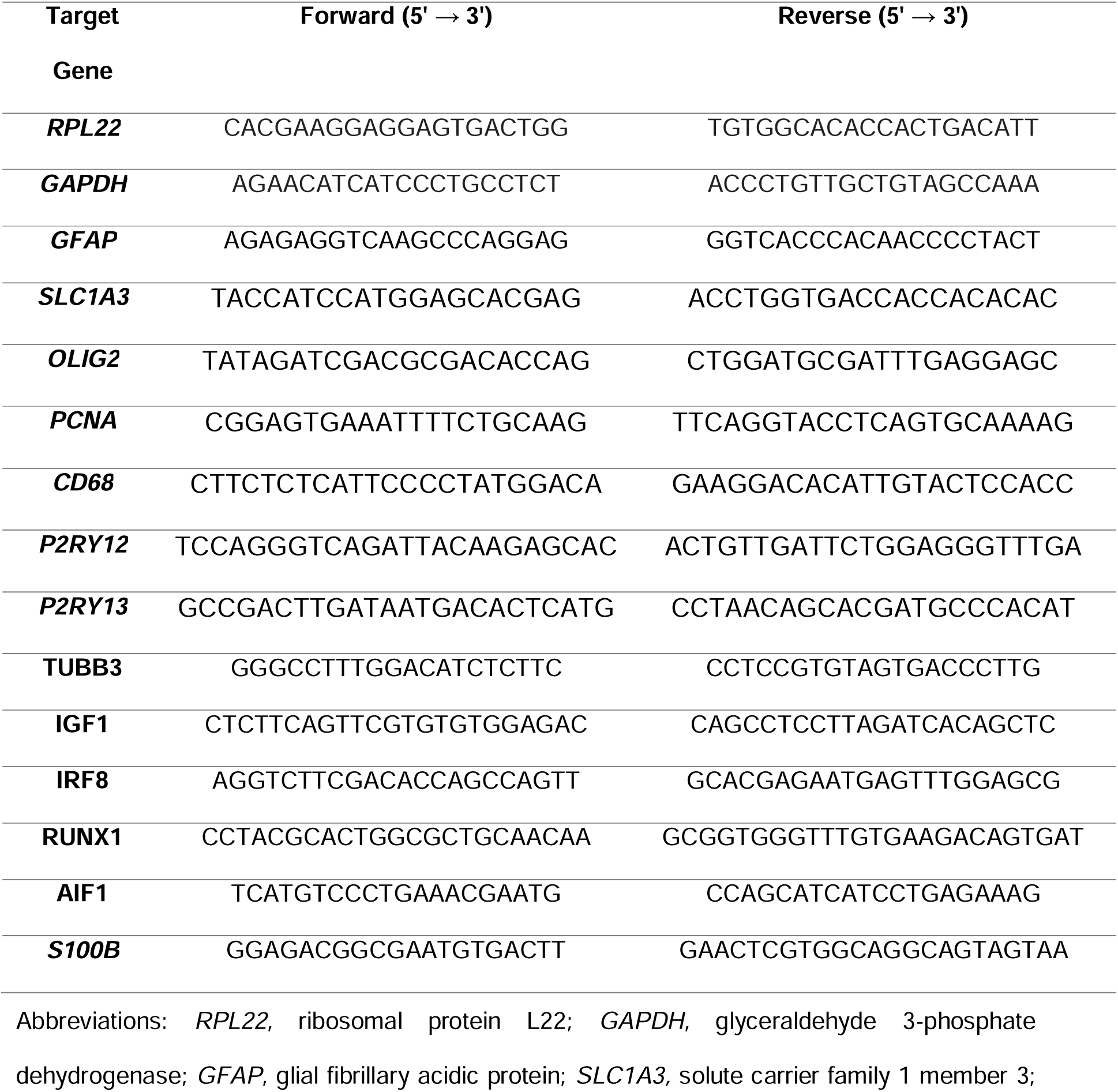
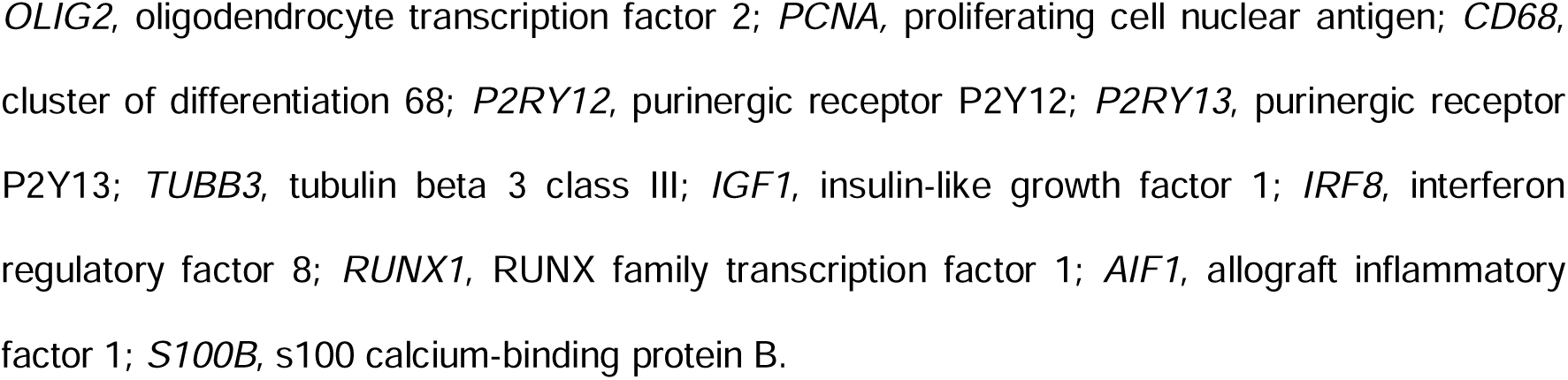
Primer sequences used for reverse transcriptase quantitative polymerase chain reaction analysis.

### Antibody arrays

Cell culture supernatants were collected at day 7 of STB culture and centrifuged at 1000 x *g* for 2 min. Supernatant from neurosphere cultures with and without iEMPs were evaluated for the presence of 40 human cytokine proteins, using an inflammation membrane antibody array, and 7 MMP and 3 TIMP proteins, using a human MMP antibody array (Abcam, Catalog #ab134003 and # ab134004, respectively) according to the manufacturer’s instructions. The spot intensity of each protein was determined by the Protein Array Analyzer for ImageJ software. Raw densitometry data was further processed by subtracting the background using negative controls in each membrane, and the positive control signals before proceeding to analysis. Additionally, all membranes were normalized against a basal medium control and represented as mean pixel density, in arbitrary units.

### Statistical analysis

Data are expressed as mean□±□standard deviation (S.D.), with statistical analyses conducted in GraphPad Prism v9.0.1 (GraphPad Software, San Diego, CA). When comparing only two experimental groups, the unpaired Student t-test was used for data with normal distribution; if otherwise, the Mann-Whitney test was used. When comparing three or more groups, a one-way analysis of variance (ANOVA) followed by the Bonferroni or Tukey post-hoc test was used for data with normal distribution. To compare different groups with two independent variables, we used a two-way ANOVA followed by the two-stage linear step-up procedure of the Benjamini– Hochberg method; statistical significance was considered for p < 0.05.

## Results

### hiPSC-derived myeloid progenitors present features of erythromyeloid progeny

Microglia arise from EMPs during primitive haematopoiesis in an extra-embryonic structure, the yolk-sac, and migrate to the developing CNS(Dermitzakis et al., 2023). EMPs undergo several developmental stages marked by modulation of gene and protein expression (Tay et al., 2017). Here, we employed a commercially available method for differentiating hiPSCs into microglia progenitors and characterised them along differentiation. Employing three different hiPSC lines (Figure 1 A), we observed morphological changes in the colonies at various time points during the hiPSC differentiation process, from elongated and adherent to round-shaped semi-adherent cells, resembling in vivo development (Supplementary Figure 1 A). On day −1, hiPSC were collected and plated at a low colony density (4-10 colony/cm^2^). Cell pluripotency was confirmed by the high detection of the TRA-1-60 and TRA-1-81 antigens (98.5 ± 0.7% and 94.4 ± 0.6%, respectively) and Stage Specific Embryo Antigen (SSEA)-4 (93.6 ± 1.9%) (Supplementary Figure 1 B), and the low expression of SSEA1 (2.4 ± 0.4%), a member of cluster of differentiation antigens (Supplementary Figure 1 B). On the following day, mesodermal patterning was induced. Intra-colony proliferation and outward migration of mesodermal cells could be observed (Supplementary Figure 1 A). Two distinct cell populations emerged: (i) non-adherent primitive iEMPs from day 6 (Supplementary Figure 1 A, white arrows) and (ii) elongated cells resembling nearby endothelium that firmly adhered to the substrate (Supplementary Figure 1 A, hollow arrows). EMP generation and development upon primitive haematopoiesis is a tightly controlled transcriptional program (Speicher et al., 2019).

**Figure 1.**
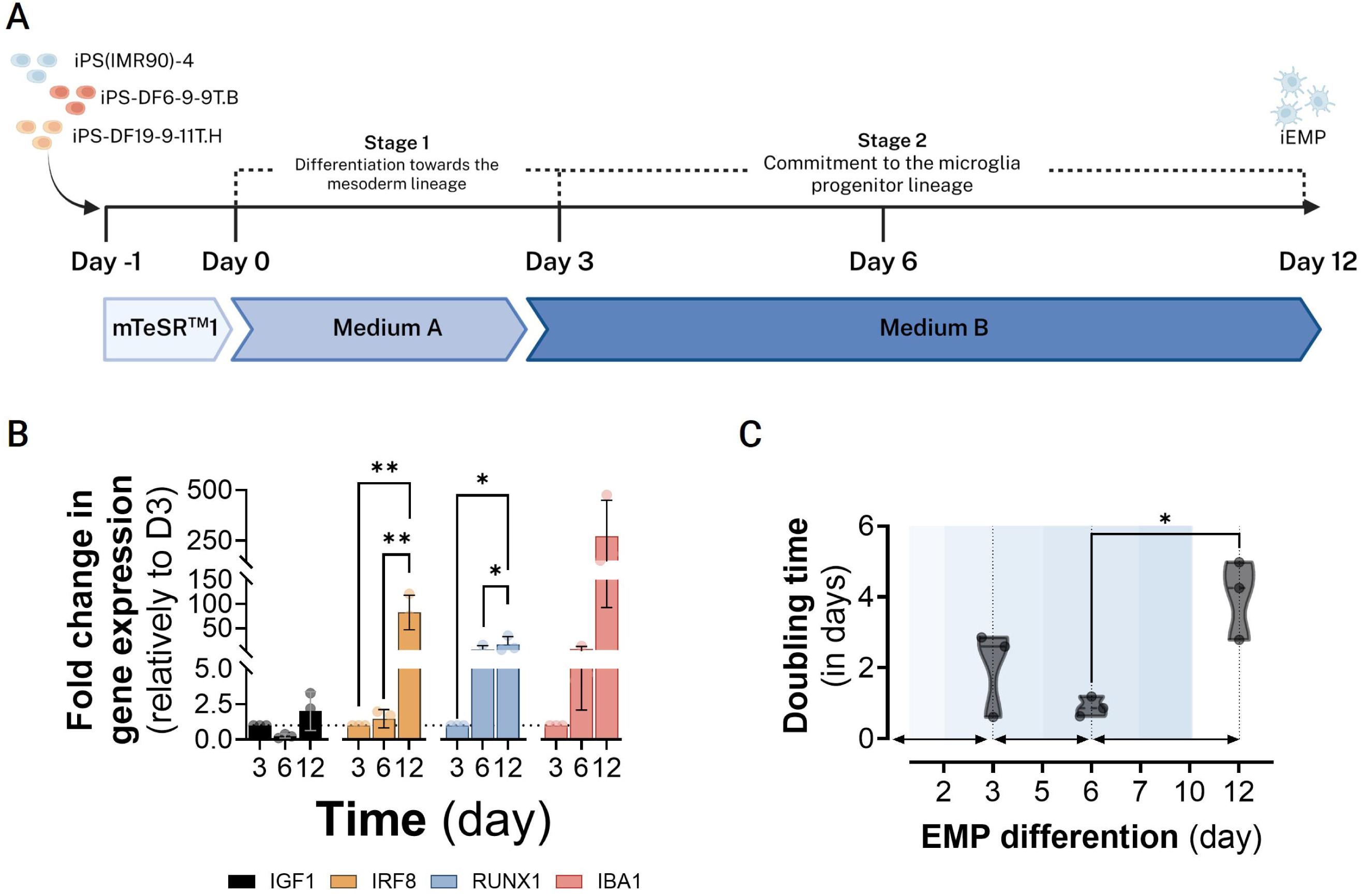
Gene modulation of EMP-associated genes throughout hiPSC differentiation in 2D. hiPSC-derived microglia progenitor cells (erythromyeloid progenitors, iEMPs) modulate EMP-specific transcription factor and microglia gene expression. Data was generated using RT-qPCR and the comparative cycle threshold values method (2 −ΔΔ*Ct*) was used. Results are represented as Δ*Ct* (target/reference ratio), being RPL22 used as a reference. Data represented as mean ± S.D., N = 3 (three hiPSC lines).

IRF8 and RUNX1 transcription factors have been described as vital for the development of this progeny. During hiPSC to iEMP differentiation, a time-dependent upregulation of EMP transcription factors was observed (Figure 1 B). IRF8, RUNX1 and AIF1 presented a 55.8-, 2.4- and 35.9-fold increase respectively, from day 6 to 12 (iEMP-specification period), concomitant with the appearance of the EMP-like cells (free-floating, Supplementary Figure 1 A). In this time period we also observed a significant 4.5-fold increase in population doubling time from day 6 to 12 (p < 0.05), suggesting an increased proliferation of the latter, although inter-cell line variability could be denoted (Figure 1 C).

On day 12, cells with typical EMP morphology were detected as round, free-floating cells in the culture supernatant (Supplementary Figure 1 A, yellow arrows). These cells were CD43-positive (96.8 ± 0.5%), and partially positive for CD34 and TREM2 (7.1 ± 0.7% and 6.8 ± 0.9% respectively), similar to progenitor cells from the primitive program of the YS (McQuade et al., 2018; Atkins et al., 2021); less than 1.0% of the population was positive for pluripotency markers (Supplementary Figure 1 C).

### hiPSC-derived EMPs (iEMPs) integrate neurospheres in 250 mL STB operated under perfusion

Aiming to develop a reliable and controllable process for producing large quantities of neurospheres containing iEMPs for preclinical research purposes, we cultured three hiPSC lines in the Ambr® 250 Modular system (Figure 2 A, B). The Ambr® 250 systems, modular and high-throughput (HT), are scalable, multi-vessel 250 mL stirred-tank bioreactor systems, which enable parallel screening. These systems have been successfully employed in process development for biopharmaceutical production (Tai et al., 2015; Roth et al., 2017; Wilson et al., 2019; Klimpel et al., 2024; Silva et al., 2024), vaccine development (Yang et al., 2023; Fang et al., 2024), and more recently, cell and gene therapy applications (Costariol et al., 2019; Rotondi et al., 2021; Müller et al., 2025), namely for the production of stem-cell products (Ho et al., 2022). The iNSCs were seeded as single cells into STB vessels operated in batch mode, to facilitate NPC interaction and neurosphere formation (Simão et al., 2016, 2018). The STB vessels used in this study are unbaffled and have a single, wide impeller geometry (elephant-ear, diameter 30 mm and 45° pitch-blade angle). This vessel design allows for improved 3D cell culture suspension, homogenization maintaining a low shear-stress (Rotondi et al., 2021; Ho et al., 2022), The initial agitation speed was set at 160 rpm based on tip speed conversion between the Ambr® 250 Modular vessels and our previous system (70 rpm) (Simão et al., 2016, 2018). Small clusters formed within 6 hours post-inoculation, and by day 2, uniform and compact neurospheres were observed amongst all cell lines, similar to our previous data in another STB system (Figure 2 C and Supplementary Figure 2 A, B). Additionally, in the first 48 hours, all cell lines proliferated at least 1.6 times showing the STB robustness for the culture of different hiPSC lines (Supplementary Figure 2 C). On day 2, the co-culture was initiated by the addition of iEMP and co-culture medium (co-culture condition); co-culture medium alone was used as control (medium control condition, MCtrl). We adjusted the agitation rate from 160 (day 0) to 230 rpm over 7 days to tailor homogeneous and size-controlled neurospheres while maintaining high cell viability, (Figure 2 C-G and Supplementary Figure 2 D). No significant differences were found in cell density, neurosphere concentration, diameter, and area between iEMP-neurosphere co-cultures and iNPCs in their expansion medium (Ctrl, Figure 1 C-G) or in the co-culture medium (MCtrl, Supplementary Figure 2 D, E). In all conditions, cell density, neurosphere diameter and area increased throughout the 7 days of culture, which are indicative of cell proliferation (Figure 2 D, F, G). The cells’ proliferative capacity was evaluated using EdU labelling to track nascent DNA (Flomerfelt and Gress, 2015). Similar levels of EdU incorporation were obtained amongst mono- and co-cultures, which was further supported by the comparable cell densities and the low percentage of CD45 and EdU double-positive cells when compared to NPCs (approximately 6% vs 27%, Figure 2 C, G, H and Supplementary Figure 2 E).

**Figure 2.**
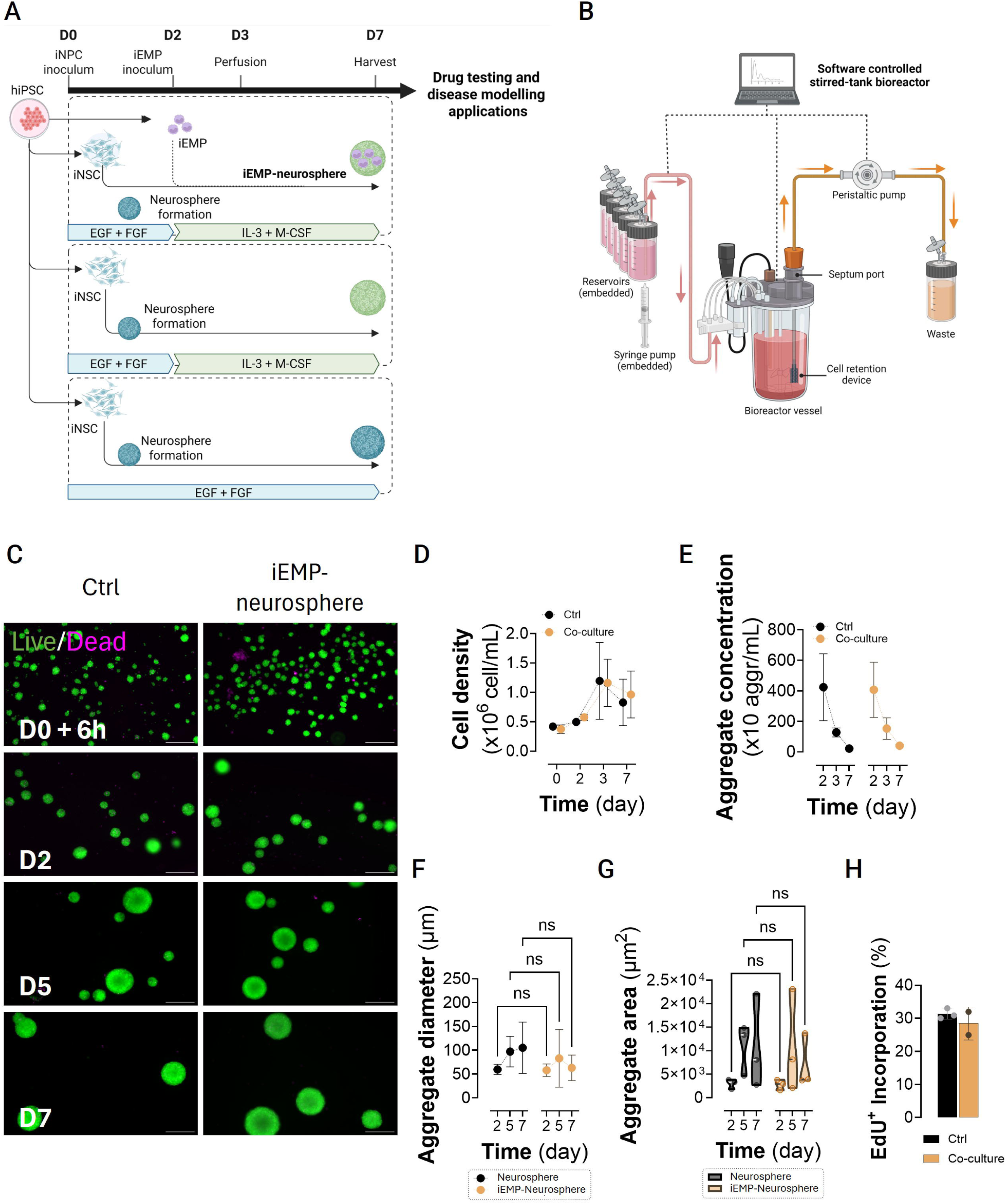
Neurosphere generation and iEMP co-culture in the single-use Ambr® 250 Modular system. Schematic representation of the (a) experimental protocol and (b) the single-use Ambr® 250 Modular system bioreactor vessel layout, including adapted perfusion system. Three STBs were inoculated with hiPSC-derived neural progenitor cells (iNPCs) at day zero and cultured for 7 days to form neurospheres or further inoculated with hiPSC-derived erythromyeloid progenitors (iEMPs) at day 2, to generate co-cultures (iEMP-neurospheres). (c) Representative live/dead images of neurospheres and iEMP-neurospheres 6 h post-inoculum, 2, 5 and 7 days of culture. Aggregates were stained with FDA (live cells, green) and PI (dead cells, red). The scale bar represents 200 *µ*m. (d) Cell density, (e) aggregate concentration, (f) diameter and (g) area measured during the 7 days of culture. Average diameter was quantified by measuring at least 100 aggregates per condition. (h) Proliferative profile of the cultured cells by incorporation and immunostaining of EdU along differentiation. Quantification of EdU incorporation was performed by counting EdU-positive and total (DAPI-positive) cells in three different iNSpheroid optical fields of three technical replicates, using the cell counter plug-in of ImageJ software. Data are shown as mean ± S.D. of three biological replicates (three hiPSC lines).

Neurosphere formation in STBs is a two-stage process, with initial cell-cell interactions and small cluster formation that occurs in the first 2 days, followed by cluster fusion and cell proliferation to form neurospheres (Bayó-Puxan et al., 2018; Simão et al., 2018). Herein, we observed the same behaviour in iEMP-neurosphere co-cultures (Figure 2 D). After an initial aggregation stage, from day 2 to 7, neurosphere concentration decreased, while the diameter increased due to neurosphere fusion and cell proliferation. Thus, the co-culture of iEMP-neurospheres did not impair the typical neurosphere aggregation process, sustaining high viability and typical neurosphere morphology.

iEMPs were co-cultured with neurospheres as a single-cell suspension, in a medium supplemented with M-CSF and IL-3, crucial proteins for maintaining EMP survival and proliferation (Douvaras et al., 2017; Haenseler et al., 2017). To maintain a constant concentration of neurotrophic and EMP factors and the removal of spent medium and cell debris, a cell retention device was adapted to the STB system (Figure 2 B). This cell retention device was adapted to fit the septum cap port of the headplate of the Ambr250 vessel and was comprised of a rubber adaptor, a metallic deep tube and a stainless-steel frit sparger (small cylindrical metallic mesh with a pore range of 10-30 μm). On day three of culture, the STBs were switched from batch to perfusion-operating mode. From that day onward, a constant metabolite concentration was observed in the co-culture, with no significant differences from the control conditions (Supplementary Figure 3). To our knowledge, these results demonstrate for the first time the feasibility of operating the Ambr250 modular system in a perfusion mode.

### iEMP differentiation mimics CNS developmental trajectory, under co-culture with iNPCs

Next, we sought to analyse the differentiation trajectories of iEMPs by investigating the expression of EMP and microglia markers: *IRF8*, *RUNX1*, *AIF1*, *CD68*, *P2RY12* and *P2RY13* after co-culture (Figure 3 A). After the introduction of iEMPs in neurosphere cultures, a tendency for the up regulation of these genes was also detected along the neurosphere expansion period (days 3 to 7, Figure 3 A). RUNX1 and lineage-specific transcription factor PU.1 were also detected in the iEMP-neurospheres by immunostaining at 5 days of co-culture (day 7 of neurosphere culture, Figure 3 B). On this day, IRF-8 and TMEM119 positive cells were also observed in the iEMP-neurospheres, indicating a microglia lineage commitment (Figure 3 C, D). Interestingly, we observed that not all iEMP infiltrated the neurospheres, with viable single cells observed in the supernatant even at 7 days of co-culture (Supplementary Figure 3 B). Immediately upon harvest from the STB, these cells started to adhere to the surface of the culture dish and acquired an elongated morphology. Upon 1-2 hours of plating, these cells presented a more ramified morphology with small protrusions and were IRF8- and TMEM119-double positive (Supplementary Figure 4 B), suggesting that the soluble factors of the co-culture support iEMP survival and differentiation as single-cell suspension.

**Figure 3.**
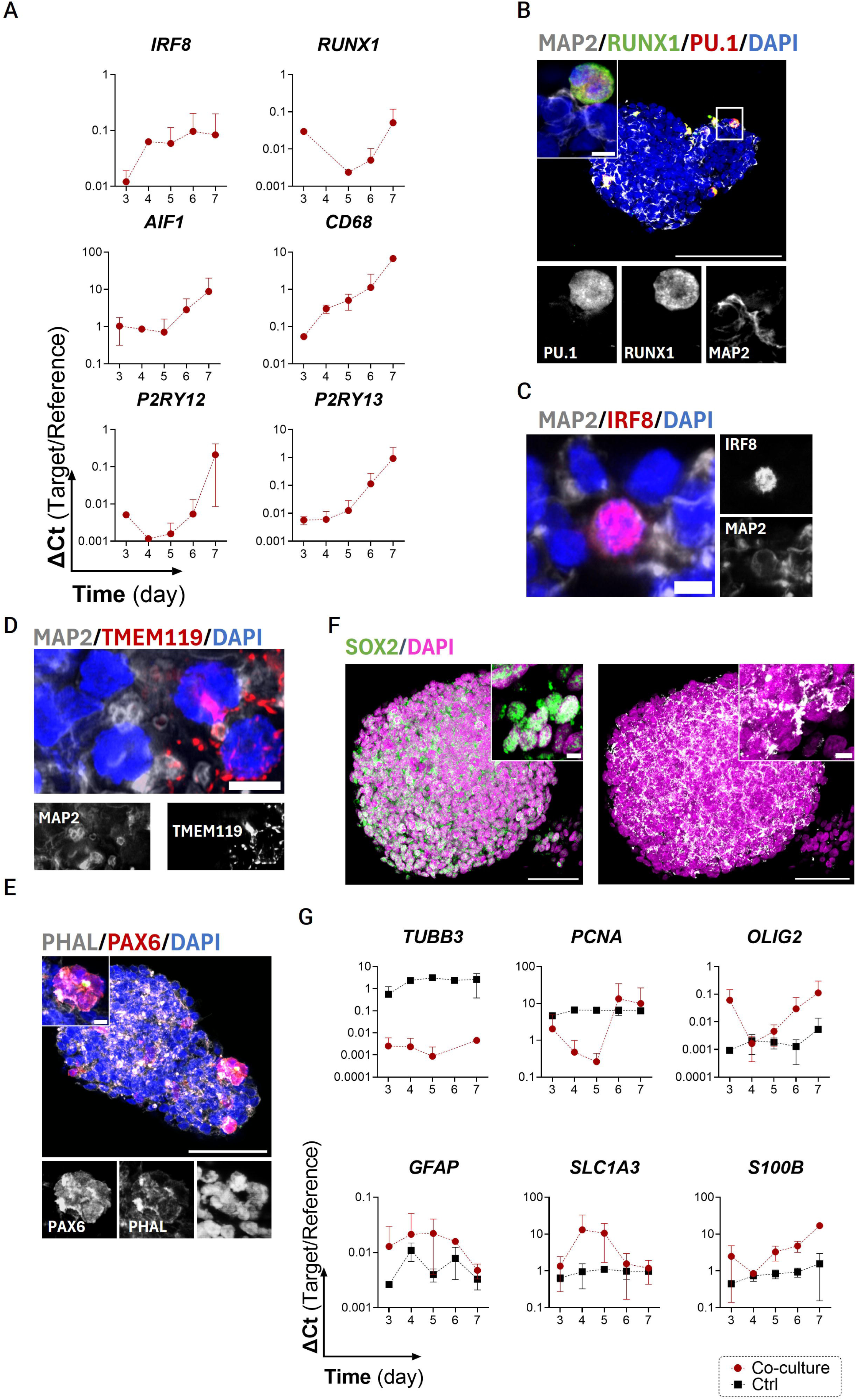
Microglia progenitors maintain an iEMP-like phenotype in co-culture with iNPCs in neurospheres. (a) Gene expression profiling of EMP-related genes in the iEMP-neurospheres along culture (days 3-7). Data was generated using RT-qPCR and the comparative cycle threshold values method (2 −ΔΔ*Ct*) was used. Results are represented as Δ*Ct* (target/reference ratio), being RPL22 used as a reference. Data represented as mean ± S.D., N = 3 (three hiPSC lines).Immunofluorescence images showing expression of (b) RUNX1 (green), PU.1 (red), and MAP2 (grey); (c) IRF8 (red) and MAP2 (grey); (d) TMEM119 (red) and MAP2 (grey); (e) phalloidin (PHAL, grey) and PAX6 (red); and (f) SOX2 (magenta); and (g), *β*III-tubulin (grey) in the neurospheres. All immunostainings were counterstained with DAPI (blue for (b)-(e), and magenta for (f)). Scale bars: 50 *µ*m (all except for (c, d, f) for which the scale bar is 5 *µ*m) with an optical slice thickness of 0.5 *µ*m. For zoom-in images, scale bar = 5 *µ*m. Images were obtained by applying maximum intensity projection of 10 optical Z-slices. (g) Gene expression analysis of neurospheres in presence or absence of iEMP. Data was generated using RT-qPCR and the comparative cycle threshold values method (2 −ΔΔ*Ct*) was used. Results are represented as Δ*Ct* (target/reference ratio), being RPL22 used as a reference. Data represented as mean ± S.D., N = 3 (three hiPSC lines).

We also addressed the impact of the co-culture on the iNPC phenotype. PAX6- and SOX2-positive neural progenitors were detected at day 7 in the neurospheres (Figure 3 E, F). The presence of β-III tubulin progenitors/immature neurons was also observed, indicative of spontaneous differentiation (Figure 3 F), which has been reported previously (Simão et al., 2018). By assessing genes for neuronal differentiation, *TUBB3*, oligodendrocyte progenitor cells, *OLIG2*, and astrocytes, *GFAP*, *SLC1A3* and *S100B*, no major differences were found in the presence or absence of iEMP, suggesting that the latter do not impact the neural progenitor’s phenotype. Taken together, these results demonstrate that iEMPs progress in the differentiation program towards the microglia fate when co-cultured with iNPC neurospheres in stirred-tank bioreactors.

### The secretome of iEMP-neurospheres is enriched in drivers of ECM remodelling and neuronal differentiation

Previous reports on NPC-microglia crosstalk have shown that the microglia secretome promotes neuronal differentiation, being enriched in cytokines and chemokines with both pro- and anti-inflammatory functions (Osman et al., 2019; Rosin et al., 2023). We investigated the secretory profile of 7-day iEMP-neurosphere co-cultures in comparison to neurosphere monocultures. Cytokine array analysis revealed significantly higher secretion of the chemoattractant proteins, CXCL1, CXCL8 and CCL2, and osteopontin, a non-collagenous bone matrix glycoprotein; and a decreased concentration of CXCL9, G-CSF and FLT3L in iEMP-neurospheres in comparison to the monoculture (Figure 4 A and Supplementary Figure 4 C-E).

**Figure 4.**
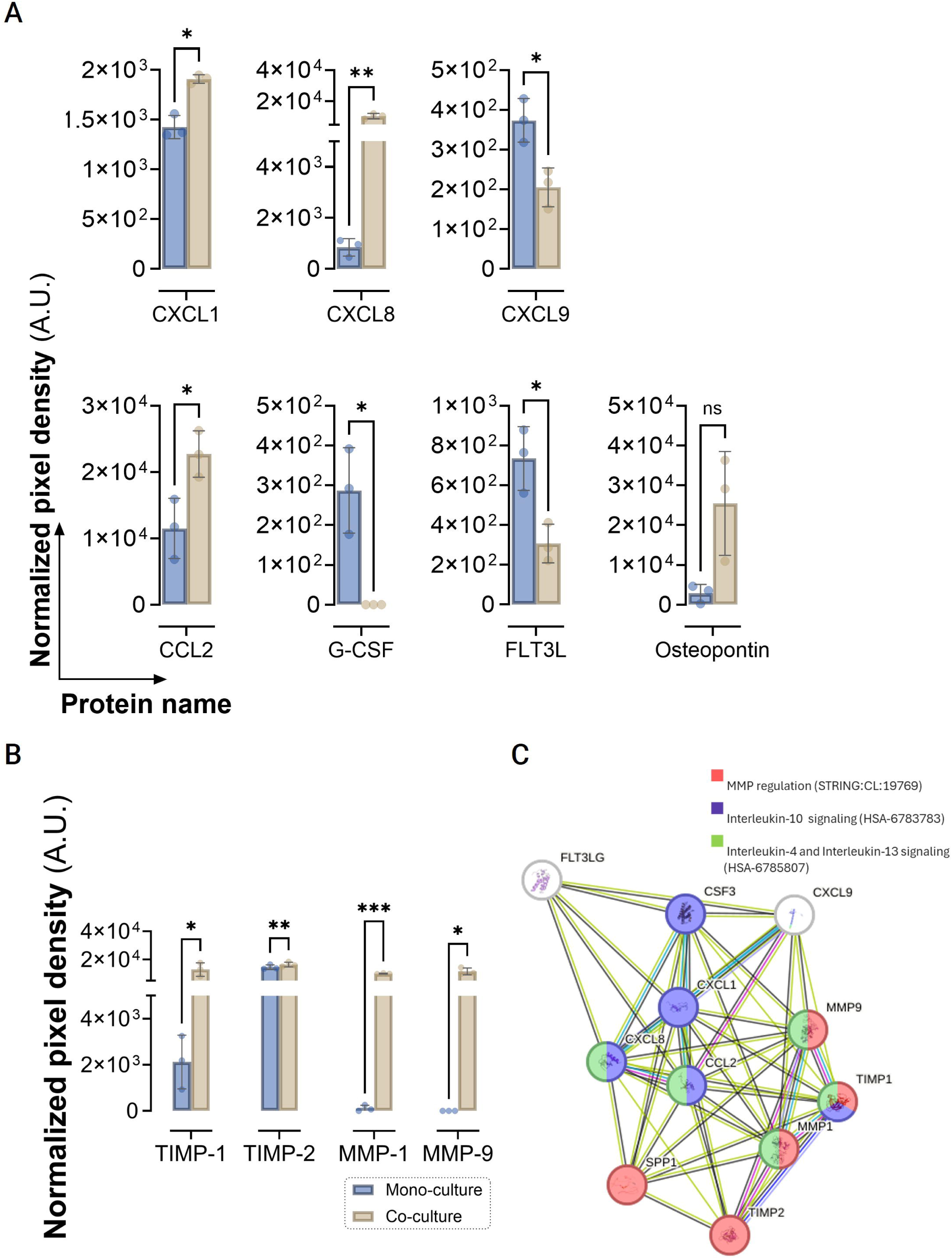
Neurosphere secretome profiling at day 7 underlines iEMP modulation of the microenvironment in comparison to mono-cultures. (a) Cytokine, chemokine and (b) extracellular matrix remodeling-related protein secretion. Data is represented as pixel density (arbitrary units, A.U.) normalized by a reference membrane. Data represented as mean ±. S.D. of 3 biological replicates (hiPSC lines). (c) STRING representation of the significantly modulated proteins. For complete data refer to Supplementary Figure 4.

MMP-8 and −9 have been reported to be highly expressed in migrating/amoeboid developing microglia during brain embryogenesis. In the iEMP-neurospheres, we found a significantly higher detection of MMP-1 and MMP-9 (Figure 4 B and Supplementary Figure 4 F, G). Concomitantly, tissue inhibitor of metalloproteinase (TIMP)-1 and −2 were significantly secreted in the co-culture condition (Figure 4 B and Supplementary Figure 4 F, G). These proteins have been reported in several physiological functions during CNS development, namely oligodendrocyte and neuronal differentiation (Pérez-Martínez and Jaworski, 2005; Sulik and Chyczewski, 2009; Nicaise et al., 2019; Ferreira et al., 2023). To investigate potential functional interactions among significantly modulated proteins identified in the co-culture secretome, a STRING network analysis was performed. The resulting protein-protein interaction network revealed distinct clusters enriched in biological processes related to MMP regulation (STRING:CL:19769, red nodes), IL-10 signaling (HSA-6783783, blue nodes), and combined IL-4 and IL-13 signaling pathways (HSA-6785807, green nodes). Central nodes in the network, such as MMP9, CXCL1, and TIMP1, highlight key regulatory hubs potentially mediating neural-iEMP crosstalk within the co-culture environment. These findings suggest an active remodeling of the extracellular matrix and cytokine signaling milieu in response to the presence of iEMP. Altogether, these results point towards an iEMP-induced modulation of the microenvironment.

## Discussion

Herein, we successfully developed a scalable and controlled system for co-culturing hiPSC-derived iEMPs and iNPCs as three-dimensional (3D) neurospheres. This model recapitulates key features of early iEMP differentiation and facilitates the study of early interactions between EMPs and NPCs.

The use of the Ambr® 250 Modular system, adapted for perfusion, represents a significant advancement for parallelizing complex hiPSC-based cultures, as the ones highlighted in this study. This system allows for precise control over environmental conditions, such as temperature, pH, and dissolved oxygen, which are critical for maintaining cell viability and function over extended culture periods (Ho et al., 2022) (reviewed in (Qian et al., 2016b)). The ability to operate in perfusion mode ensures a continuous supply of nutrients and removal of waste products, thereby supporting long-term cultures and reducing operator intervention, enhancing overall reproducibility among batches. Indeed, we observed that by adapting a stainless-steel frit sparger as a cell retention device that is incorporated in the bioreactor vessel through the septum cap, we were able to maintain metabolite concentration throughout the neurosphere cell proliferation. Cell density, viability and neurosphere formation were similar amongst batches and hiPSC lines used, demonstrating the robustness of such STB system which is essential for improving overall cell quality. Additionally, the use of this scalable bioreactor system allowed for parallelization of multiple conditions and cell donors, reaching faster results with enhanced reproducibility and higher experimental throughput.

Microglia progenitors arise early during embryonic development from an extraembryonic structure called the yolk sac (Prinz et al., 2019). A first wave of myeloid cell development takes place between E7.0 and E8.0 in a process known as primitive haematopoiesis that leads to the generation of EMP cells. EMP cells give rise to yolk sac subpopulation, A1 (cKit^+^RUNX1^+^CD45^+^CX3CR1^-^) cells, followed by A2/amoeboid (RUNX1^+^PU.1^+^CD45^+^CX3CR1^+^MMP-8/-9^+^) cells (Prinz and Priller, 2014; Tay et al., 2016; Prinz et al., 2019). The latter will further differentiate into embryonic microglia, which will mature in the developing CNS into ramified microglia.

The biological relevance of the iEMP-neurosphere model is underscored by recapitulating some morphological traits and gene and protein expression features to migrating microglia progenitor cells (amoeboid EMP). Our results demonstrate that iEMPs in co-culture with neurospheres express key transcription factors such as RUNX1 and PU.1, as well as IBA-1, indicating a microglial lineage commitment. The secretion of matrix metalloproteinases (MMPs), particularly MMP-9, further supports the notion that these cells are adopting a phenotype similar to embryonic microglia. MMPs play crucial roles in CNS development, including neuronal migration, myelination, and synaptic plasticity (Yong et al., 2001; Agrawal et al., 2008). MMP-9 plays a crucial role in sensory circuit development during early postnatal critical periods by regulating synaptogenesis, axonal pathfinding, and myelination (reviewed in (Reinhard et al., 2015)). Its activity is tightly controlled through enzymatic degradation and inhibition by thrombospondins or TIMP-1, which is co-secreted in response to synaptic activity and can form a complex with MMP-9 before secretion (Roderfeld et al., 2007). Interestingly, our data revealed the concomitant secretion of MMP-9 and TIMP-1 in the co-culture of iEMP-neurospheres in comparison to the neurospheres alone. This might suggest that iEMP have an impact in the co-culture secretome towards a neurogenic effect.

The secretome analysis further highlighted the secretion of CXCL1, CXCL8 and CCL2 in the iEMP-neurosphere co-culture. Although typically considered as effector cytokines during inflammation, with known functions in microglia and astrocytic activation and blood brain barrier permeability (Ashutosh et al., 2011; He et al., 2016; Dubový et al., 2018; Joly-Amado et al., 2020; Michael et al., 2020), some literature has suggested that these chemoattractants have roles in physiological neurodevelopmental processes. CCL2/CCR2 network has been linked to an increased potential for dopaminergic neuronal differentiation (Edman et al., 2008) and cell excitability, dopamine release, and locomotor activity in rats (Guyon et al., 2009). Moreover, CCL2 has been identified at different embryonic stages in relation to the cytoarchitectural organization of the CNS, implying a role for this chemokine during brain development (Zhen Meng et al., 1999; Rezaie et al., 2002). CXCL1 and CXCL8 have been linked with progenitor cell division, neurogenesis, and tyrosine hydroxylase(TH)-positive cell number in rodent precursor and neurosphere cultures (Edman et al., 2008).

This study comprises a first step towards models with innately differentiated microglia. Variability between cell lines in terms of genetic stability, differentiation potential, and cell morphology remains a challenge in hiPSC-based research (Volpato and Webber, 2020). This variability can affect the differentiation potential and functionality of the derived cells. In this work, we have used three different hiPSC lines that varied in donor gender, cell origin (somatic versus PBMC) and method of reprogramming (viral and non-viral based). The robustness and reproducibility shown for the neurosphere formation, proliferation and differentiation across different hiPSC lines are notable and indicate the potential of employing this co-culture strategy for further disease modelling and drug screening applications, using the Ambr® 250 Modular system.

In conclusion, the system we developed offers a powerful platform for studying CNS cell interactions in a controlled and physiologically relevant environment. The ability to precisely control and modulate environmental conditions in this 3D model enables detailed investigations into key aspects of neurodevelopment and disease, offering significant potential for preclinical research. Future studies leveraging this platform can deepen our understanding of microglia in both health and disease, potentially guiding the development of new therapeutic strategies targeting microglial dysfunction.

## Author Contributions

Conceptualization, C.M.G. and C.B.; methodology, C.M.G. and C.B.; Investigation, C.M.G., I.Q.S., M.D.; writing—original draft, C.M.G. and C.B.; writing—review & editing, C.M.G, I.Q.S, M.D., P.M.A., C.B; funding acquisition, C.B., P.M.A; resources, C.B., P.M.A; supervision, C.B.

## Acknowledgements

The authors acknowledge Dr. Daniel Simão and Dr. Giacomo Domenici for useful discussion in building the scientific hypothesis.

Illustrations were created in biorender.com.

---

**Supplementary Figure 1 - Phenotypic characterization of hiPSC-EMP differentiation protocol.**

(a) Schematic representation of the iEMP differentiation protocol using three different hiPSC lines. (b) Representative bright-field imaging of iEMP differentiation, along the 12-day protocol. (c) Quantification of the cell population expressing pluripotency-associated proteins within hiPSC collected at day −1 of the differentiation. Data was collected by flow cytometry analysis of hiPSC suspension, from the three cell lines under analysis. (d) Cell doubling time at days 3, 6 and 12 of differentiation. (e) Quantification of the cell population expressing EMP-associated proteins within the iEMP collected at day −1 of the differentiation. Data was collected by flow cytometry analysis of a iEMP cell suspension, from the three cell lines under analysis. All data graphs are represented as mean ± S.D.

**Supplementary Figure 2 - Bioprocess characterization during the first seven days of STB culture.**

Representative live/dead images from day 2 neurospheres produced in the (a) DASGIP® Parallel Bioreactor System (hiPSC-1) and in (b) Ambr® 250 Modular system (representation of hiPSC-1, −3 and −4). Neurospheres were stained with fluorescein diacetate (FDA, live cells, green) and propidium iodide (PI, dead cells, red). Scale bar, 200 *µ*m. (c) Cell density at the inoculum (0) and after 2 days of culture, in hiPSC-1, −3 and −4. Representation of N > 2 for each cell line. Average fold increase of day 2 versus day 0 is plotted for each cell line. (d) Representative live/dead images of medium control (MCtrl) after 6hours, 2-, 5- and 7-days post-inoculum. Neurospheres were stained with fluorescein diacetate (FDA, live cells, green) and propidium iodide (PI, dead cells, red). Scale bar, 100 *µ*m. (e) Percentage of actively proliferating cells in culture along culture time, assessed by incorporation of EdU.

**Supplementary Figure 3 - Adaptation of a perfusion system to the Ambr® 250 Modular system bioreactor vessels.**

GLC, AQB, GLN, PYR, *NH*3, LAC and LDH specific concentration in the culture supernatant of the different conditions. Data represented as mean ± SD of three independent experiments (three different hiPSC lines).

**Supplementary Figure 4 - iEMP gene expression and protein secretion profile after seven days of co-culture with the neurospheres.**

(a) Representative live/dead (upper panel) and phase contrast (lower panel) images of iEMP-neurospheres after 7-days of culture. Neurospheres were stained with fluorescein diacetate (FDA, live cells, green) and propidium iodide (PI, dead cells, red). Scale bar, 200 *µ*m. (b) Immunofluorescence images showing expression of phalloidin (PHAL), IRF8 (green), and TMEM119 (red) of iEMP collected from the supernantant of the STB cultures, after 7 days. Cells were allowed to adhere for 1-2 hours in PLOL-coated coverslips. Scalebar 50 *µ*m. (c-e) Extracellular matrix remodeling, and (f,g) cytokine and chemokine-related protein secretion. Representative antibody array membranes showing detection of (c) 40 cytokines and chemokines, (d) 80 cytokines and chemokines, and (f) MMP and TIMP proteins under control and co-culture conditions after 5 days in culture. All examples shown (c, d, f) are from iPSC line DF6-9-9T.B.Data is represented as pixel density (arbitrary units, A.U.) normalized by a reference membrane. Data represented as mean ±. S.D. of 3 biological replicates (hiPSC lines). For comparing three or more groups two-way ANOVA test with Šídák’s multiple comparison test was used: (*) p < 0.05; (**) p < 0.01; (***) p < 0.001; (****) p < 0.0001.

