## Supplementary figures and images for "Engineering a perfusion bioreactor system for hiPSC-derived progenitor co-culture capturing microglial features in CNS development"

### Fig. S1

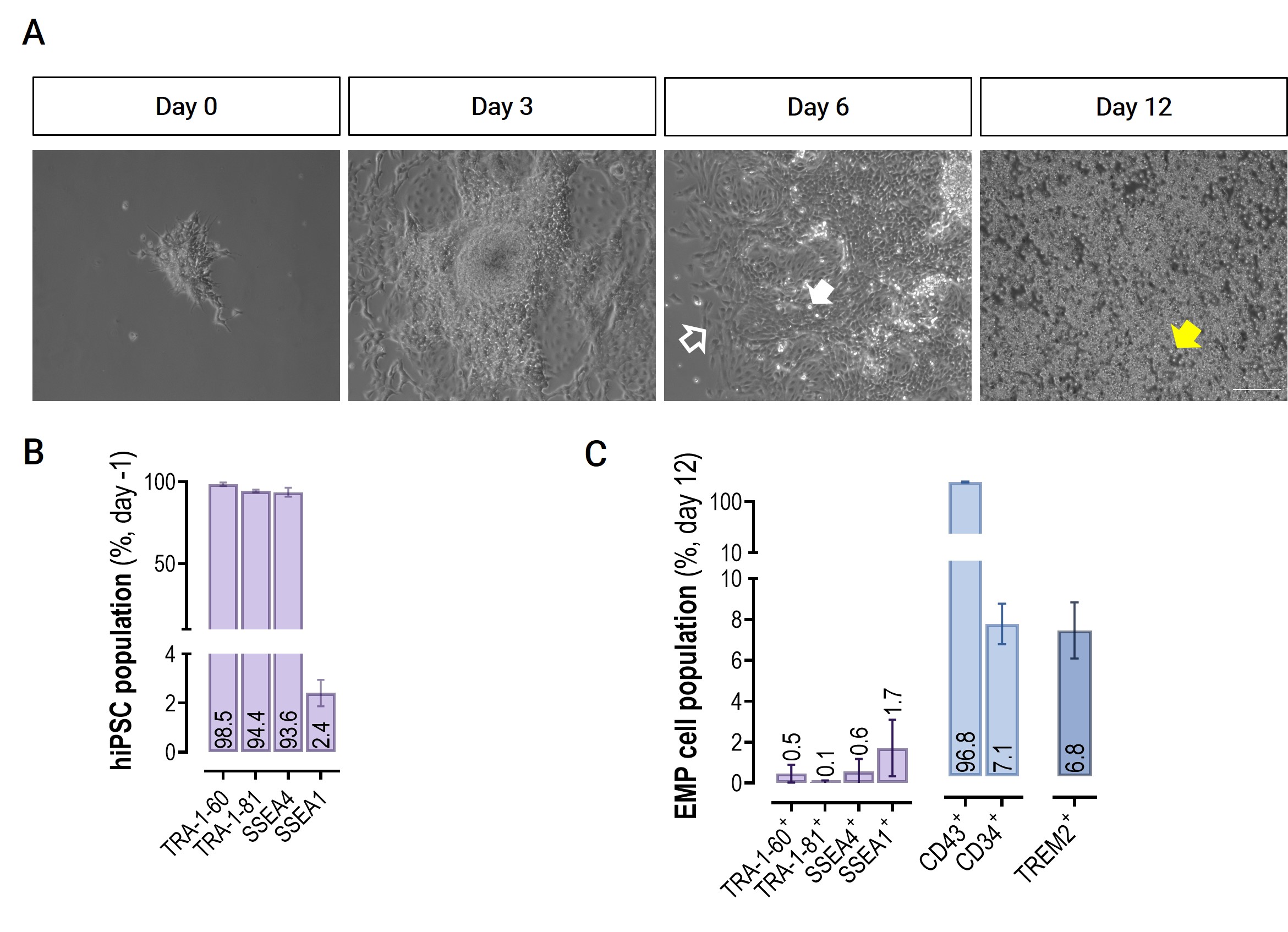

### Fig. S2

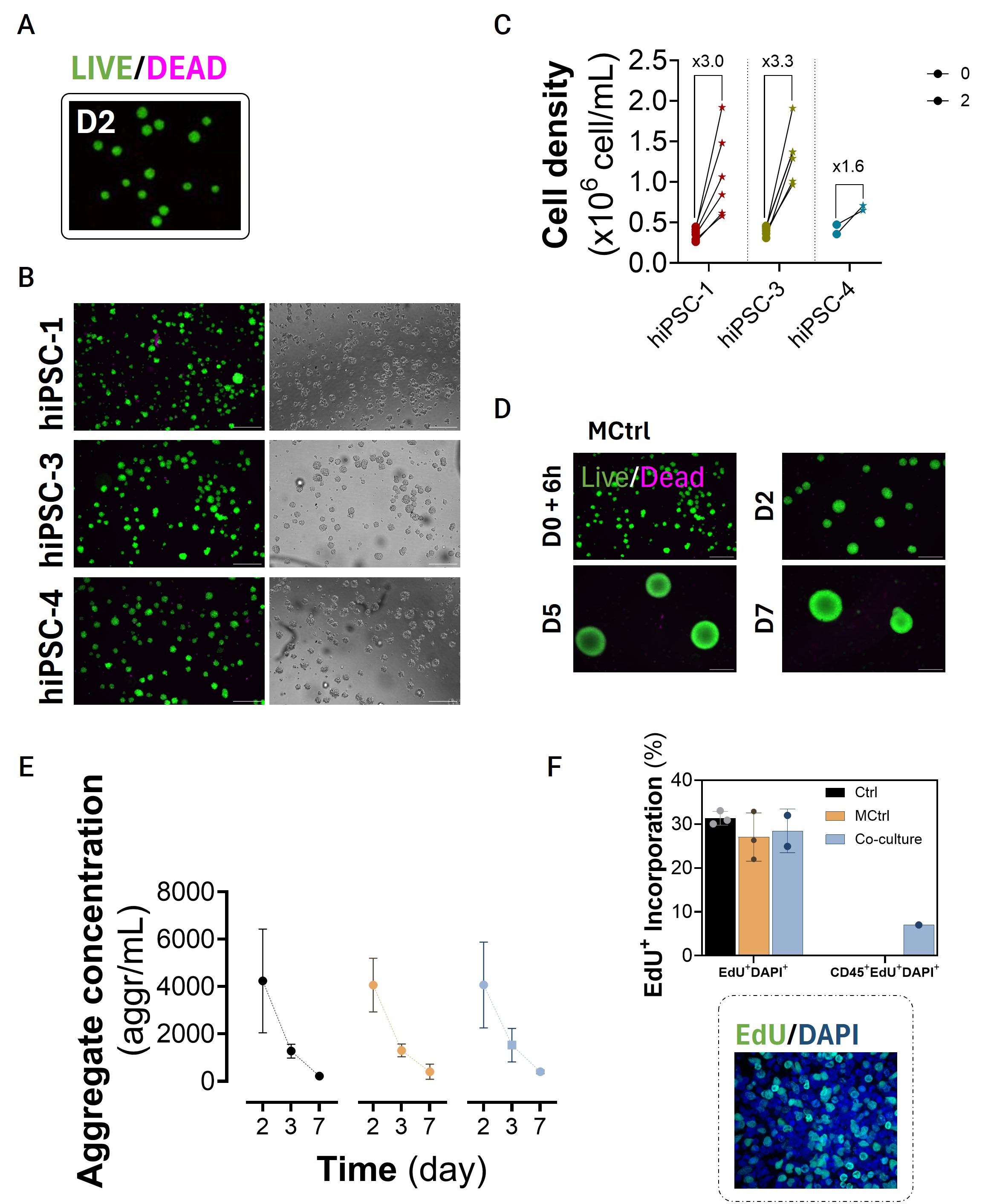

### Fig. S3

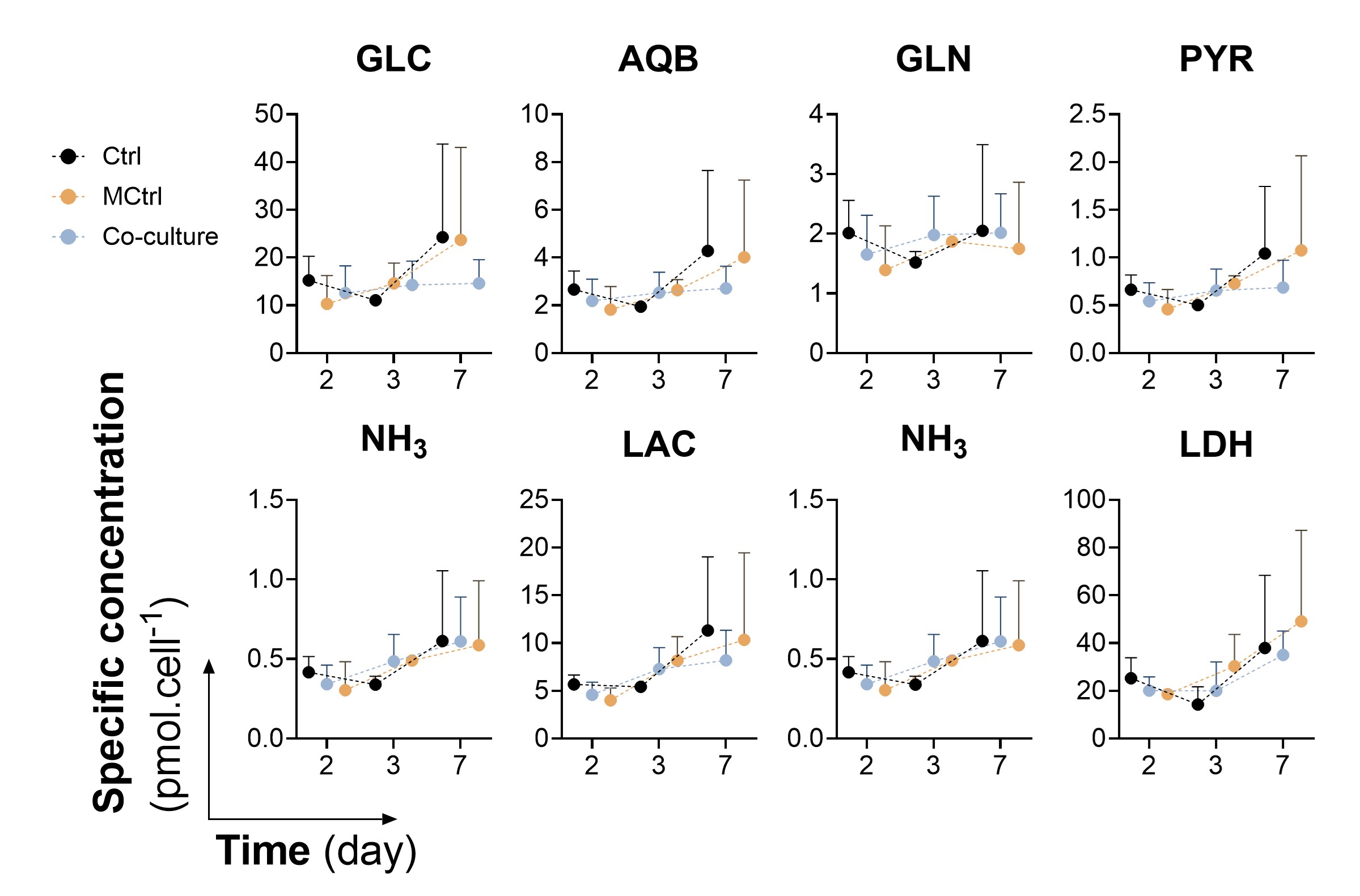

### Fig. S4

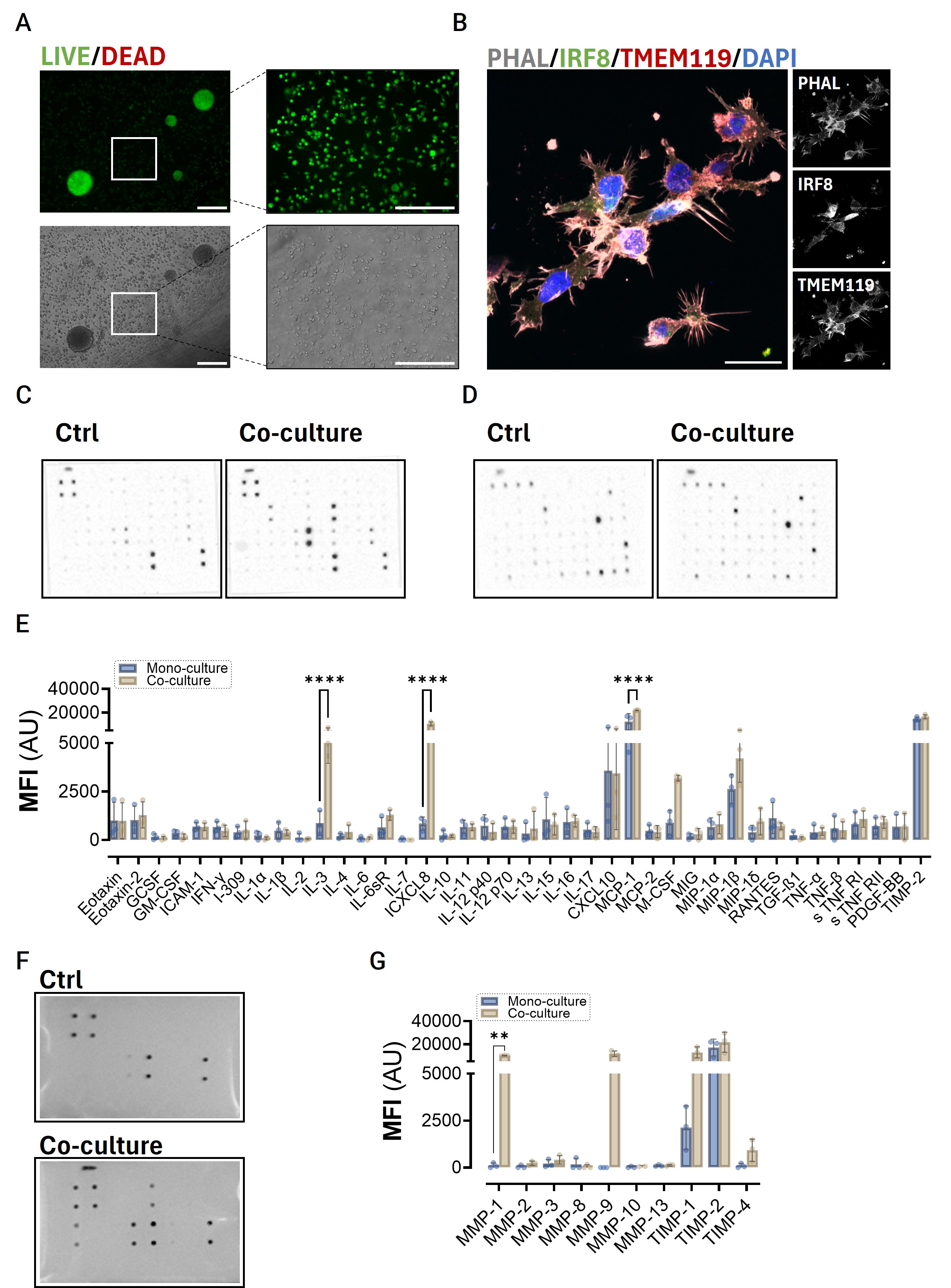
